# Enhanced reflexive eye closure is mediated by ipRGC signals and is specific to migraine with visual aura

**DOI:** 10.1101/2020.12.12.422528

**Authors:** Eric A Kaiser, Harrison McAdams, Aleksandra Igdalova, Edda B Haggerty, Brett Cucchiara, David H Brainard, Geoffrey K Aguirre

## Abstract

**Objective:** To assess the contribution of the melanopsin-containing, intrinsically photosensitive retinal ganglion cells (ipRGCs) and the cones to reflexive eye closure as an implicit measure of interictal photophobia in migraine.

**Methods:** We studied twenty participants in each of three groups: headache-free (HAf) controls, migraine without aura (MwoA), and migraine with visual aura (MwA). Participants viewed spectral pulses that selectively targeted melanopsin, the cones, or their combination while we recorded orbicularis oculi electromyography (OO-EMG) and blinking rate.

**Results:** Time course analysis of OO-EMG demonstrated that reflexive eye closure was tightly coupled to the spectral pulses. Compared to both the MwoA and HAf control groups, the MwA group had enhanced OO-EMG and blinking activity in response to melanopsin and cone stimulation in combination and in isolation. This response scaled with the contrast of the stimulus.

**Conclusions:** Our findings suggest that ipRGC signals, whether elicited by melanopsin stimulation or from presumed extrinsic cone input, provide the afferent input for light-induced reflexive eye closure in a photophobic state. Participants with migraine and visual aura had a distinctly different response to visual stimulation as compared to the other two groups. This is in contrast to prior findings in this same cohort in whom higher explicit ratings of visual discomfort were found for both MwA and MwoA as compared to controls. Such a dissociation suggests distinct pathophysiology in forms of migraine, interacting with separate neural pathways by which ipRGC signals elicit implicit and explicit signs of visual discomfort.

## Introduction

People with migraine experience light sensitivity during a headache,^1-4^ as well as photophobia between headaches (i.e., interictally).^5-9^ While often assessed by self-report, reflexive eye closure in response to light may also be used as an index of visual discomfort,^10, 11^ and appears to be implemented by a sub-cortical reflex arc independent of the visual cortex.^12^ Rodent studies suggest that the afferent arm of light aversion begins with signals from the melanopsin-containing, intrinsically photosensitive retinal ganglion cells (ipRGCs).^13-16^ As ipRGCs also receive input from the cone photoreceptors, both melanopsin and cone signals conveyed by the ipRGCs may contribute to light aversion (Figure 1a).

**Figure 1.**
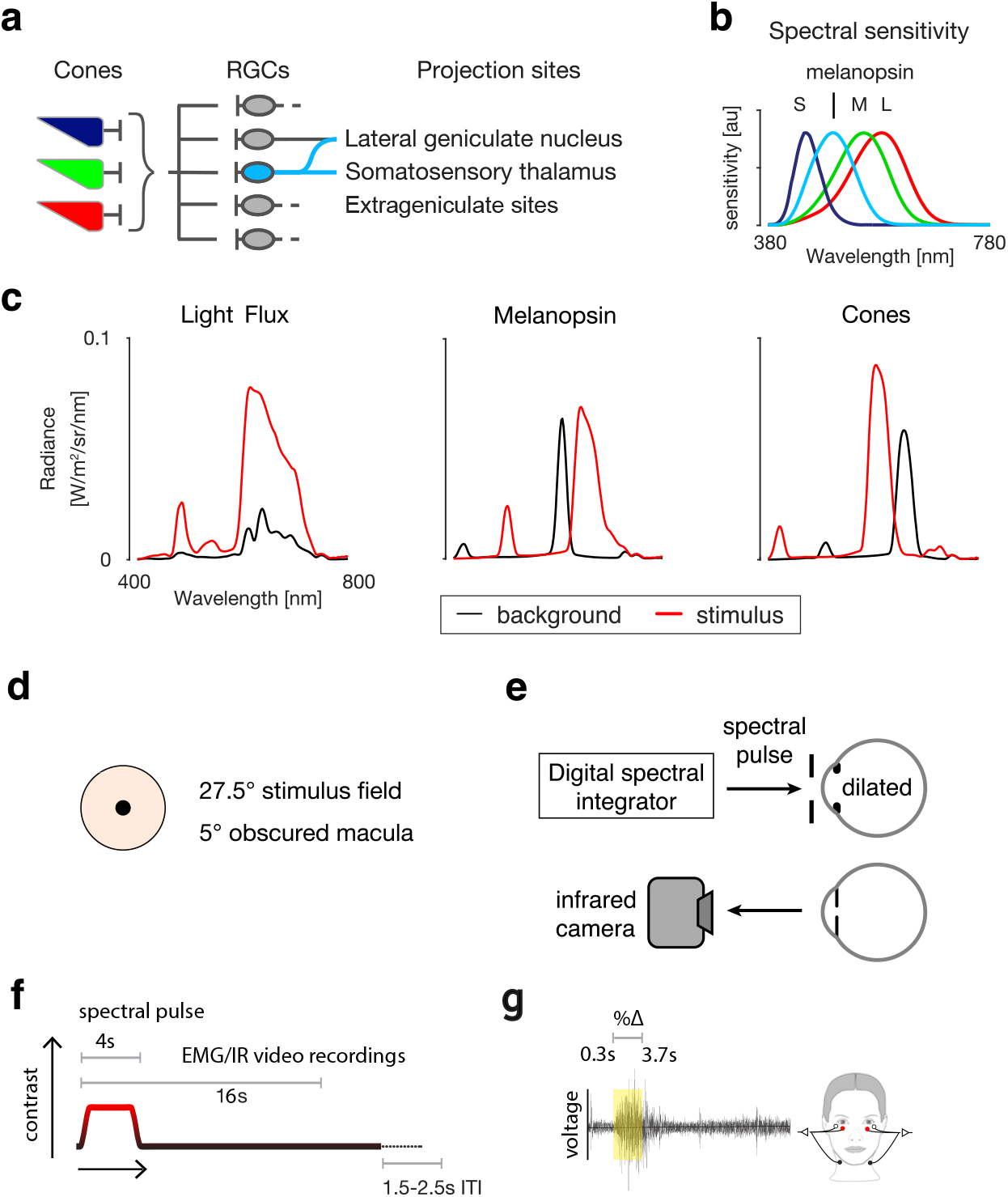
Experimental Overview. **a**. There are multiple classes of melanopsin-containing ipRGCs that vary in their central projections, function, and extent to which they receive input from cones. The ipRGCs project to the lateral geniculate nucleus, somatosensory thalamus and extrageniculate sites including the brainstem and hypothalamus. **b**. The spectral sensitivity functions of the relevant photoreceptors under daylight conditions. **c**. Shown are pairs of spectra (background: black; stimulus: red) that differ in excitation for the targeted photoreceptors. From left to right, the stimuli produce: equal contrast on the cones and melanopsin (termed light flux); contrast only on melanopsin; and equal contrast across all three classes of cones but no contrast on melanopsin. **d**. The stimulus spectra were presented in an eyepiece with a 27.5-degree diameter field, with the central 5 degrees obscured to minimize macular stimulation. **e**. The light from a digital spectral integrator was presented to the pharmacologically dilated right eye of the participant through an artificial pupil. Blinking of the left eye was recorded with an infrared (IR) camera. To record orbicular oculi electromyography (OO-EMG), electrodes were placed inferior to both eyes. **f**. Each trial featured a four-second pulse during which the stimulus transitioned from the background to the stimulation spectrum and back. EMG and IR camera recordings started prior to the onset of the pulse and continued for 16 sec. There was an inter-trial interval that varied between 1.5 and 2.5 s. **g**. OO-EMG activity was calculated from the percent change in the standard deviation of the voltage from baseline over a 3400 msec window starting 300 msec after pulse onset and ending 300 msec before pulse offset.

Explicit report of visual discomfort in people with migraine and interictal photophobia appears to result from a selective amplification of ipRCG signals, after the integration of melanopsin and cone inputs.^17^ Here we ask if the same findings are present for two implicit measures of visual discomfort: electromyography of the orbicularis oculi (OO-EMG) and blink frequency. These were recorded while participants observed spectral pulses that selectively targeted the cones and/or melanopsin (Figure 1b, 1c). We studied twenty participants in each of three groups: headache-free (HAf) controls, migraine without aura (MwoA), and migraine with visual aura (MwA). We find that both melanopsin and cone stimulation, in isolation and in combination, lead to reflexive eye closure. This effect is enhanced in the MwA group, compared to both the MwoA group and the HAf controls.

## Methods

In a pre-registered study, we used silent substitution stimulation to selectively target melanopsin, the cones, or both. We presented 4-second pulses of these stimuli while we recorded OO-EMG, and blinking in the contralateral eye via an infrared camera. The participants and stimuli presented here have been the subject of a prior paper on pupil responses and self-reported visual discomfort;^17^ the current data were collected in the same experimental sessions.

### Participants

A total of 60 participants (ages of 25 to 40) were recruited from Philadelphia and the University of Pennsylvania. All recruited participants underwent screening using the Penn Online Evaluation of Migraine:^18^ an automated headache classification using the International Classification of Headache Disorders (ICHD)-3-beta criteria. It also incorporates a set of published questions regarding ictal and interictal photophobia that were scored with a point for each “yes” response to questions 1 through 7 (referred to here as the Choi score).^19^ Recruited participants also completed the Visual Discomfort Score (VDS) survey;^20^ our scoring has been previously described.^17^ Candidate participants were required to meet all inclusion criteria for one of three groups:

1. Migraine with visual aura (MwA): a) classification of migraine with visual aura by the POEM, b) Choi score of 6 or 7, c) a response of “yes” to the Choi query regarding the presence of photophobia during headache-free periods, d) one or more headaches within the prior month.
2. Migraine without aura (MwoA): a) classification of migraine without aura by the POEM, b) Choi score of 6 or 7, c) a response of “yes” to the Choi query regarding the presence of photophobia during headache-free periods, d) one or more headaches within the prior month.
3. Control: a) classification of mild non-migrainous headache or headache-free by the POEM, b) a response of “No” or “I don’t know” to a question regarding a family history of migraine, c) a response of “No” to a question regarding a history of childhood motion sickness, d) VDS score of 7 or lower.

Exclusion criteria related to impaired vision, inability to collect usable pupillometry, head trauma, and seizure history have been previously described.^17^ Participants were not excluded based on medication use and were allowed to continue to take their current medications during data collection.

### Stimuli

We designed stimuli that target specific photoreceptor classes through silent substitution.^21^ In this approach, sets of light spectra are created that have the nominal property of producing equal excitation of one or more “silenced” classes of photoreceptors and varying excitation on one or more “targeted” photoreceptors. We targeted three main photoreceptor mechanisms: melanopsin, the cones, or their combination (referred to as “light flux”) (Figure 1b, c).

Stimuli were generated as described in prior reports.^17, 22, 23^ Briefly, we used a digital light synthesis engine (OneLight Spectra, Vancouver, BC, Canada) that produces stimulus spectra under digital control at 256 Hz. We created separate background and stimulation spectra that provided 1) a nominal 400% unipolar Weber contrast on melanopsin while silencing the cones (melanopsin-directed background/stimulus pair), 2) 400% contrast on each L-, M-, and S-cone classes while silencing melanopsin (cone-directed background/stimulus pair), and 3) 400% contrast each on melanopsin and each L-, M-, and S-cone classes (light flux background/stimulus pair) (Figure 1c). Background spectra for each stimulus type differed in luminance but had similar chromaticity.^17^ We also produced contrast levels of 100 and 200% for each stimulus direction by scaling the relevant stimulus spectra. We did not explicitly silence rods or penumbral cones,^22^ although we believe that the properties of our stimuli minimize the contribution of these photoreceptors.

Stimuli were presented through a custom-made eyepiece with a circular, spatially uniform field of 27.5° diameter. The central 5° (diameter) of the field was obscured to minimize effects of the foveal macular pigment (Figure 1d). Apparatus calibration and stimulus validation have been described previously.^17^

### Experiment Structure

Participants were studied during multiple sessions, usually held on different days. To reduce variation in circadian cycle across sessions, subsequent sessions were initiated within three hours of the time of day when that participant started their first session.

Acclimation to the testing room and apparatus and pharmacologic dilation of the right eye has been previously reported.^17^ Afterwards, the participant remained in darkness for the remainder of the experiment. Participants viewed the stimuli through their pharmacologically dilated right eye and a 6 mm diameter artificial pupil to control retinal irradiance.

On each trial, the participant viewed a pulsed spectral modulation designed to target melanopsin, the cones, or both (Figure 1c). The transition from the background to the stimulation spectrum (melanopsin, cones, or light flux), and the subsequent return to the background, were windowed with a 500-msec half-cosine. This minimized the entopic percept of a Purkinje tree in the melanopsin-directed stimulus.^22^ The duration of the pulse was 4 seconds, after which the stimulus field returned to the background spectrum (Figure 1d). EMG and infrared camera recordings continued for another twelve seconds after the pulse ended. After each trial, there was a variable inter-trial-interval of 1.5 – 2.5 seconds (uniformly distributed) that reduced the predictability of the onset of the next trial.

Ten consecutive trials that targeted the same photoreceptor direction but varied in contrast were grouped together into an acquisition. The ordering of the contrast levels (100, 200, 400%) was counterbalanced,^24^ the first trial was discarded so that all retained trials had controlled first-order stimulus history. A total of 6 acquisitions, 2 of each stimulus type, comprised a single session. Acquisitions were ordered such that consecutive acquisitions were not of the same stimulus class. We attempted to gather 4 sessions of data for each participant but retained all participants who completed two sessions that contained at least six acceptable trials—as judged by pupillometry—for each stimulus type. Participants did not complete all 4 sessions for a variety of reasons.^17^

### Electromyography

We recorded OO-EMG starting prior to pulse onset and ending 12 seconds after pulse offset (Figure 1e,f). This measure is sensitive to tonic (squinting) and phasic (blinking) eye closure. There was a 1.1-sec delay between the start of the trial and the initiation of EMG recording due to a slow computational operation; thus, the period of EMG recording prior to pulse onset was approximately 400 msecs. Prior to electrode placement, the skin surface was wiped with an alcohol pad. Two small reference electrode pads were placed inferior to each eye, and a ground lead was placed on the neck. EMG was recorded with the BioNomadix 2-Channel EMG equipment (Biopac Systems, Inc). Participants were informed that the electrodes were measuring “eye responses”.

### Blink quantification via infrared videography

We recorded blinks from the left eye of the participant (contralateral to the eye receiving stimuli) using an infrared (IR) camera (Pupil Labs GmbH) mounted on a post ∼25 mm from the eye. The camera has two IR LEDs mounted adjacent to the lens, providing illumination of the eye. A 60-Hz video clip was recorded for each trial, starting 1.5-s prior to pulse onset and ending 12-s after pulse offset (Figure 1e,f). These videos were processed using custom software (https://github.com/gkaguirrelab/transparentTrack).^25^ Blinking was quantified as the number of video frames in which the glints (first Purkinje images) from the active IR light sources were absent.

### Analysis

Data analysis was performed using custom MATLAB code (Mathworks). EMG activity was quantified by calculating the standard deviation of the recorded voltage within a sliding 500-msec window. The activity time series were normalized as percentage change relative to activity occurring prior to pulse onset.

First we examined the time-course of EMG responses for each group of participants, averaged across the three stimulus types (cone, melanopsin, and light-flux). The mean OO-EMG response across trials and stimuli at each contrast level was calculated, and then averaged across participants.

We quantified the effect of our stimuli upon the OO-EMG and blink measure by calculating the average response over a 3400-msec window starting 300 msecs after pulse onset and ending 300 msecs prior to pulse offset. This window was selected to avoid the influence of blinking at stimulus onset or offset in the measured response.^26^ Our pre-registration had proposed a 4000-msec window, shifted 1000 msecs after stimulus onset; we report the results using this window in the supplementary materials. For both OO-EMG activity and proportion of blink frames from the IR camera, we took the mean response across trials within a participant, and the mean across participants within a group.

Following our pre-registration proposal, we examined the statistical effect of group and stimulus type (cone, melanopsin, light flux) upon the responses obtained for the 400% contrast stimuli. We performed a 3-by-3 mixed-effects ANOVA with a between-participants factor of group and within-participants factor of stimulus type. Significant effects were examined in post-hoc testing using the Tukey procedure including group effects collapsed over the three stimulus types and group x stimulus interactions. To test the effect of contrast upon the responses, we calculated the mean across-subject response at each log-spaced contrast level for a given group and stimulus type. Then we measured the slope for these three response values and obtained the 95% CI around that slope by bootstrapping across subjects.

### Pre-Registration and availability of data and analysis code

This study was the subject of an initial pre-registration document (https://osf.io/5ry67/) and subsequent addenda (project summary page: https://osf.io/qjxdf/). We have previously summarized our deviations from these protocols.^17^ There are four additional deviations pertaining to this study: 1) switching from root mean square to standard deviation to calculate EMG activity; 2) using a shorter (3400-msec) window for the quantification of responses; 3) using a mixed-effects ANOVA across groups and stimulus directions (as opposed to a t-test of only the melanopsin stimulus); and 4) adding the quantification of the blink frames as a secondary measure. The analysis code and data are available (https://github.com/gkaguirrelab/melSquintAnalysis).

## Results

### Participant demographic and clinical characteristics

We studied 20 individuals in each of three groups: migraine with aura (MwA), migraine without aura (MwoA), and headache-free (HAf) controls. The groups were well matched for age (Table 1). The non-photophobic HAf group contained fewer female participants than the photophobic migraine groups, reflecting the higher female to male ratio observed in migraine.^27^ Participants in both migraine groups reported similar headache frequency (∼4 headache days per month) and disability from migraine, indicating that both migraine groups contained individuals with episodic migraine and moderate disability. Medication use among the two migraine groups was similar although more MwA participants reported aspirin/acetaminophen/caffeine (Excedrin™) and Triptan.

**Table 1.**
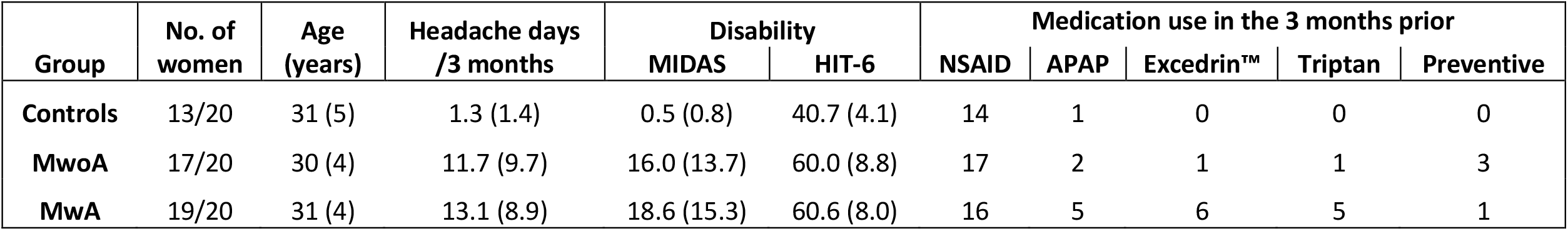
Subject demographic and clinical characteristics. Participants were asked to report the number of headache days they had experienced over the prior three months. The Migraine Disability Assessment Test (MIDAS) and the Headache Impact Test (HIT-6) measure headache disability. Where appropriate, the mean value (and standard deviation) across subjects is reported. Medication use is summarized within five categories: NSAID – non-steroidal anti-inflammatory for any indication; APAP – acetaminophen for any indication; Excedrin™ – use of any one of multiple formulations that combine acetaminophen with caffeine, aspirin, diphenhydramine, and/or phenylephrine; triptan – any 5HT_1B/D_ receptor agonist used to abort a migraine; preventive – any one of several classes of medications used to decrease headache frequency (e.g., topiramate, tricyclic anti-depressants).

| Group | No. of women | Age (years) | Headache days /3 months | Disability |  | Medication use in the 3 months prior |  |  |  |  |
| --- | --- | --- | --- | --- | --- | --- | --- | --- | --- | --- |
|  |  |  |  | MIDAS | HIT-6 | NSAID | APAP | Excedrin™ | Triptan | Preventive |
| **Controls** | 13/20 | 31 (5) | 1.3 (1.4) | 0.5 (0.8) | 40.7 (4.1) | 14 | 1 | 0 | 0 | 0 |
| **MwoA** | 17/20 | 30 (4) | 11.7 (9.7) | 16.0 (13.7) | 60.0 (8.8) | 17 | 2 | 1 | 1 | 3 |
| **MwA** | 19/20 | 31 (4) | 13.1 (8.9) | 18.6 (15.3) | 60.6 (8.0) | 16 | 5 | 6 | 5 | 1 |

### Temporal relationship of EMG activity during light stimulation trials

Each of the 60 participants viewed pulses of spectral change (Figure 1c) that targeted melanopsin, the cones, or both photoreceptor classes in combination (referred to as light flux). For each of these stimulus types, three different contrast levels were presented (100%, 200%, or 400%). Bilateral OO-EMG was recorded in participants (Figure 1g).

First, we examined the effect of the light stimulus pulse upon OO-EMG activity as a function of time across the 16-second recording interval, averaged across stimulus types within groups. For the 100% contrast stimuli, the temporal evolution of EMG activity is indistinguishable between the studied groups (Figure 2, left). There is a small increase in the EMG signal during the stimulus presentation, followed by a transient increase at stimulus offset.

**Figure 2.**
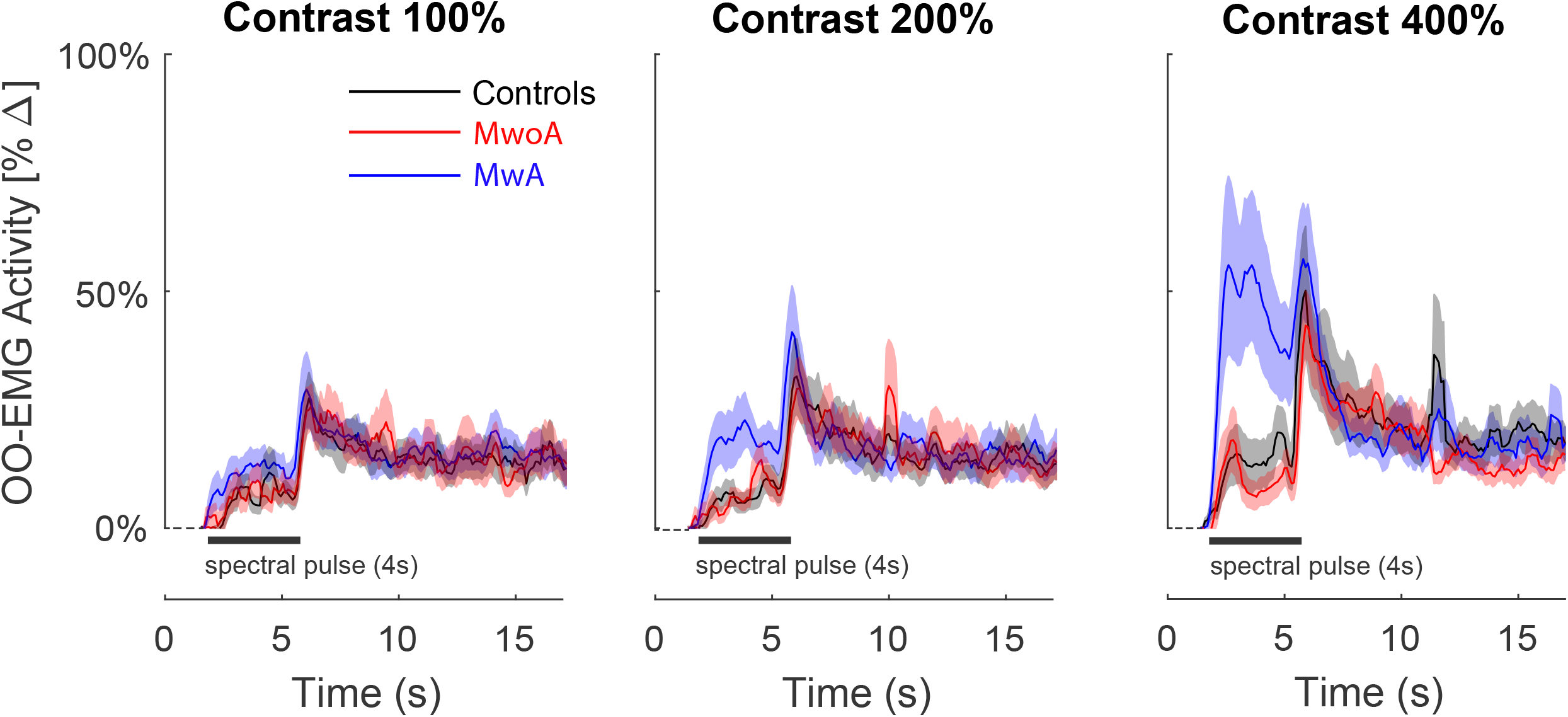
Time course of OO-EMG activity. The average OO-EMG across participants within each group (n = 20 participants per group) is shown at each contrast level (columns: 100, 200, and 400%). As we did not observe a significant difference in OO-EMG responses to light flux, melanopsin, or cone stimuli, the OO-EMG responses for the three stimulus types were averaged across stimulus types within each group. For each contrast level, responses from the three groups are superimposed and shown as a function of time over 16 second recording interval with the presentation of the 4-sec spectral pulse is labeled by a bar. The shaded area is the SEM across participants within a group. OO-EMG activity was quantified by calculating the standard deviation of voltage activity within a 500 msec window, sliding across time, which is expressed as percentage change [%Δ] relative to activity occurring prior to pulse onset.

Based on inspection of video recording of the eye contralateral to the stimulus, we interpret the offset response as participants suppressing blinks during the stimulus and engaging in increased blinking following stimulus offset. The response to the 200% contrast stimulus is similar (Figure 2, center), with the exception of an increased EMG response during the spectral pulse in only the MwA group. This effect becomes pronounced in response to the maximal, 400% contrast pulse, for which only the MwA group demonstrates a large change in EMG activity during the stimulus (Figure 2, right).

### Melanopsin and cone contrast elicits increased OO-EMG activity in MwA

We next quantified the change in OO-EMG activity during the stimulus pulse for each group, contrast level, and stimulus type. We compared OO-EMG activity from the pre-stimulus baseline to activity during the middle 3400 msec of the stimulus pulse. This response window is shorter than the 4000-msec window proposed in our pre-registration document and was selected after examination of the data shown in Figure 2, to avoid the non-specific, transient blinking response that we observed at stimulus offset.

In MwA participants, light flux pulses elicited elevated OO-EMG activity with increasing levels of contrast. This was particularly evident in response to 400% contrast which evoked an increase in mean EMG activity of 46.2% (Figure 3, right-top). Light flux evoked smaller elevations in OO-EMG activity compared to baseline in controls or MwoA participants, even at 400% contrast (mean responses: HAf controls: 17.7%, MwoA: 11.7%) (Figure 3, left-top and center-top, respectively). We also quantified the effect of contrast upon OO-EMG activity in each group, and confirmed a significant, and far larger, effect in the MwA group (Supplementary Table 1).

**Figure 3.**
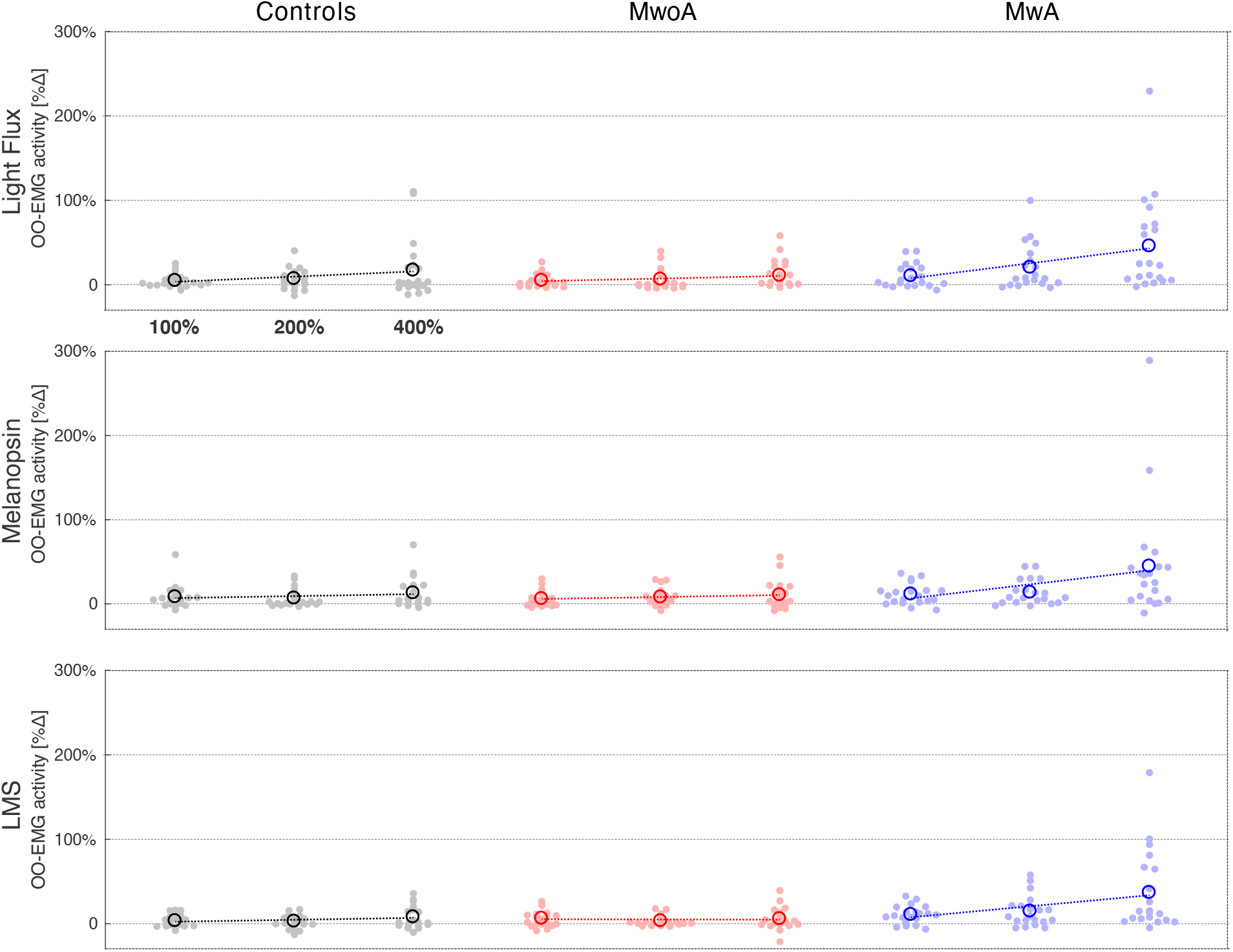
Orbicularis oculi electromyography (OO-EMG) activity by stimulus and group per post-hoc analysis window. Each row presents the OO-EMG activity evoked by stimuli that targeted a particular combination of photoreceptors, and each column contains the data from each individual group (n = 20 participants per group). The stimuli were presented at three different contrast levels (100, 200, and 400%), and these (log-spaced) values define the x-axis of each subplot. The mean (across trial) OO-EMG activity for a given stimulus and contrast is shown for each participant (filled circle), as is the OO-EMG activity across participants (open circle). The best fit line to the mean OO-EMG activity across participants as a function of log contrast is shown in each subplot. OO-EMG activity over time was quantified by calculating the standard deviation of voltage activity within a 500 msec window, sliding across time, which is expressed as percentage change [%Δ] relative to activity occurring prior to pulse onset. From these normalized time series, we calculated the average evoked response over a 3400-msec window starting 300 msec after pulse onset and ending 300 msec before pulse offset.

To assess whether the enhanced, light-induced reflexive eye closure in MwA was due to melanopsin or cone signaling, we measured OO-EMG activity in response to stimuli that targeted these two photoreceptor classes in isolation. Pulses that targeted melanopsin elicited elevated OO-EMG activity at 400% contrast (mean response: 45.1%; Figure 2, center-middle), as did 400% contrast pulses that targeted the cones (mean response: 37.2%; Figure 3, center-bottom). The OO-EMG activity in response to these stimuli was less in the HAf and MwoA groups. Similar to light flux, increasing melanopsin and cone contrast produced greater OO-EMG activity in the MwA group, and much smaller or absent effects in the other groups (Supplemental Table 1).

We examined a mixed-effects-ANOVA for the OO-EMG response to 400% contrast stimuli and found a significant effect of group (F[2,57] = 5.4, p = 0.007). The MwA group demonstrated greater EMG activity across all three types of stimuli as compared to MwoA (p = 0.011; post-hoc comparison) and HAf (p = 0.025; post-hoc comparison) (Table 2). This larger response in the MwA group was found for each stimulus direction but only a subset of responses reached statistical significance after correction for multiple comparisons (Supplemental Table 2). This mixed-effects ANOVA also showed a main effect of stimulus direction (F[2,57] = 3.5, p = 0.032), but post-hoc analyses comparing stimulus direction by group were not significant, suggesting that these effects did not differ markedly for spectral pulses targeting cones, melanopsin, or their combination.

**Table 2.** Post-hoc analyses of group effects in orbicularis oculi EMG activity at 400% contrast. We examined the effect of group upon the mean OO-EMG activity using a 3-by-3 mixed effects ANOVA. This analysis was conducted over a 3400-msec window starting 0.3 second after pulse onset and ending 0.3 second before pulse offset, which was briefer than our pre-registered analysis. This smaller window was designed to avoid observed onset and offset responses. Measured EMG activity across stimulus types (melanopsin, cones, and light flux) were averaged within subjects. The independent variables were by group (MwA, MwoA, and control) as a between subjects factor. Post-hoc testing was conducted using the Tukey procedure. *p-value < 0.05.

| Group comparison | Mean difference | t-statistic | p-value |
| --- | --- | --- | --- |
| MwA - Controls | 29.6 | 2.691 | 0.025* |
| MwA - MwoA | 33.0 | 2.995 | 0.011* |
| MwoA - Controls | 3.4 | 0.305 | 0.950 |

In our pre-registration, we proposed to quantify OO-EMG activity during a 4000-msec window, shifted 1 second from stimulus onset. When the data are analyzed using this temporal window, we find similar results, albeit somewhat attenuated by the incorporation of the non-specific blinking response at stimulus offset (Supplementary Figure 1, Supplementary Tables 3-4).

Overall, these findings indicate that both melanopsin and cone signals contribute to light-induced reflexive eye closure, and that this effect is seen primarily in the MwA group.

### Melanopsin and cone contrast elicits increased blinking in MwA

People engage in both blinking and squinting in response to a bright light. The OO-EMG signal reflects both types of muscle activity. We sought to confirm our OO-EMG findings by examining a second measure of reflexive eyelid closure. We obtained an estimate of the proportion of time spent blinking in the period following pulses of light. This was measured by identifying video frames in which the corneal “glint” of an active IR light source was obscured by the closed eyelid contralateral to the stimulus.

In MwA participants, light flux as well as targeted melanopsin and cone pulses increased the proportion of blink frames during the middle 3400 msecs of the stimulus pulse with increasing levels of contrast, particularly at 400% contrast (mean proportion of blink frames, light flux: 0.105; melanopsin: 0.092; cones: 0.086; Figure 4, right column). Lower rates of blinking were seen in the MwoA (mean proportion of blink frames, light flux; 0.048; melanopsin: 0.033; cones: 0.038; Figure 4, center column) and HAf controls (mean proportion of blink frames, light flux: 0.036; melanopsin: 0.041; cones: 0.038; Figure 4, left column). These findings parallel our measurements of OO-EMG activity. Analysis of blinking activity in relation to contrast level showed positive slopes for MwA for each stimulus type, and markedly smaller or absent effects for the other groups (Supplemental Table 6).

**Figure 4.**
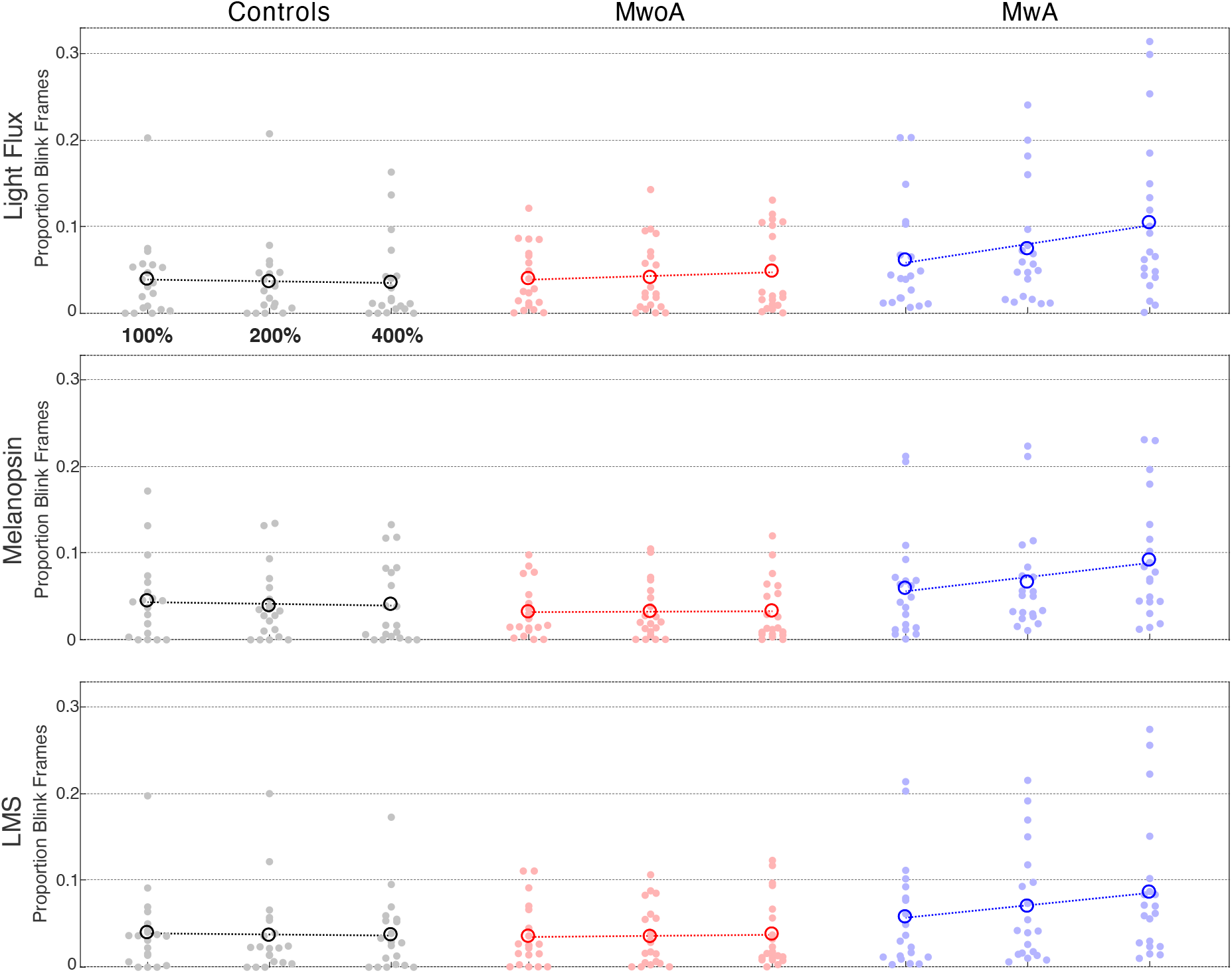
Blink activity by stimulus and group per post-hoc analysis window. Each row presents the blink activity evoked by stimuli that targeted a particular combination of photoreceptors, and each column contains the data from each individual group (n = 20 participants per group). The stimuli were presented at three different contrast levels (100, 200, and 400%), and these (log-spaced) values define the x-axis of each subplot. The mean (across trial) blink activity for a given stimulus and contrast is shown for each participant (filled circle), as is the blink activity across participants (open circle). The best fit line to the mean blink activity across participants as a function of log contrast is shown in each subplot. Blink activity is expressed as the proportion of blink frames over a 3400-msec window starting 300 msec after pulse onset and ending 300 msec before pulse offset..

A mixed-effects-ANOVA confirmed a group effect upon the blink rate in response to the 400% contrast stimuli (F[2,57] = 6.3, p = 0.003). Similar to the OO-EMG activity, the MwA group demonstrated greater blinking across all three types of stimuli as compared to the MwoA (p = 0.010, post-hoc comparison) and HAf controls (p = 0.008, post-hoc comparison) (Table 3). Post-hoc analyses confirmed that blinking activity was significantly greater in MwA participants for each of the stimuli directions at 400% (Supplemental Table 6). The mixed-effects ANOVA indicated an omnibus effect of stimulus (F[2,57] = 3.4; p = 0.038) for blinking activity. Post-hoc analyses indicated that the blinking activity was greater with light flux stimulation as compared to cone stimulation in the MwA group (p = 0.003); otherwise, all other comparisons were not significant (Supplemental Table 7).

**Table 3.** Post-hoc analyses of group effects in blink activity at 400% contrast. We examined the effect of group upon the mean proportion of blink frames using a 3-by-3 mixed effects ANOVA. This analysis was conducted over a 3400-msec window starting 0.3 second after pulse onset and ending 0.3 second before pulse offset, which was briefer than our pre-registered analysis. This smaller window was designed to avoid observed onset and offset responses. Responses across stimulus types (melanopsin, cones, and light flux) were averaged within subjects. The independent variables were by group (MwA, MwoA, and control) as a between subjects factor. Post-hoc testing was conducted using the Tukey procedure. *p-value < 0.05.

| Group comparison | Mean difference | t-statistic | p-value |
| --- | --- | --- | --- |
| MwA - Controls | 0.0562 | 3.115 | 0.008* |
| MwA - MwoA | 0.0546 | 3.025 | 0.010* |
| MwoA - Controls | 0.0016 | -0.090 | 0.996 |

Using the pre-registered 4000-msec analysis window, shifted 1 second from stimulus onset, we also observe similar results with the blinking activity data, but again find they are attenuated due to inclusion of the non-specific blinking response at stimulus offset (Supplementary Figure 2, Supplementary Tables 8-10).

Therefore, we find confirmatory evidence in a second measure that reflexive eyelid closure in response to light stimuli is greater in MwA as compared to MwoA and HAf control groups.

## Discussion

We find that both cone and melanopsin stimulation elicit reflexive eye closure. This finding implicates the ipRGCs, which both contain melanopsin and receive extrinsic input from the cones. The effect of light upon eye closure is heightened in migraine participants who have visual aura (MwA) as compared to those with migraine without aura or without headache. This is in contrast to our prior finding that, regardless of a history of visual aura, people with migraine report heightened visual discomfort to these same stimuli.^17^

### Photoreceptor contributions

We measured the degree of OO-EMG activity produced by pulses that targeted the cones, melanopsin, or their combination, at three different contrast levels. These stimuli evoke a significantly larger OO-EMG response in the MwA group as compared to the MwoA and headache-free control groups. Furthermore, OO-EMG activity increases with contrast in the MwA group in response to all three stimulus types.

In an earlier study with this cohort,^17^ we applied a quantitative, two-stage model to measurements of verbal discomfort and pupil size to characterize the relative contribution of melanopsin and cone stimulation to the responses. The model describes a non-linear combination of melanopsin and cone signals using a Minkowski metric. We are unable to apply this prior model to the current dataset. For the MwoA and control groups, there is essentially no effect of contrast to model. For the MwA group, the combination of melanopsin and cone stimulation (light flux) produces no more response than does either stimulation alone. This circumstance corresponds to an infinite Minkowski metric (winner-takes-all), which does not lend itself to a parametric characterization. A better understanding of the relative contribution of these photoreceptor inputs to reflexive eye closure might be pursued in future studies by examining stimuli that mix varying levels of cone and melanopsin contrast.

Here we examined group differences using a mixed-effects ANOVA. We find that MwA group differs from the MwoA and HAf control groups in nearly all circumstances for OO-EMG activity. The claim of a distinct response in the MwA group is buttressed by the clear temporal association of the enhanced response to the stimulus pulse, and the complementary finding of increased blinking as assessed by video analysis.

### Explicit vs. implicit measures

In surveys, people with migraine report more visual symptoms and discomfort than headache-free controls,^9, 28-31^ and these visual symptoms reduce reported quality of life.^32^ Photophobia has also been measured as the threshold of light intensity at which participants report discomfort or pain. People with migraine consistently have lower light thresholds for discomfort as compared to non-migraine controls.^5-9^ When MwoA and MwA groups have been compared, similar reports of visual discomfort and threshold light sensitivity have been found.^5, 6, 28^ In our prior study,^17^ we described the result of asking participants to verbally report their degree of discomfort to the same pulsed-stimuli studied here. We found that both MwoA and MwA reported equivalent, increased discomfort as compared to HAf controls.

Here we examined reflexive eye closure, which has previously been used as an implicit measure of visual discomfort.^10^ Reflexive blinking and squinting to bright light has been termed “dazzle” or “photic blink”, and is implemented by a sub-cortical (pre-tectal) reflex arc that does not require the visual cortex.^12^ Our current study finds that responses from this brainstem reflex arc are altered primarily in people with MwA. A recent study found that people with migraine respond with a threshold amount of OO-EMG activity at lower levels of light compared to healthy controls; however, this study did not distinguish between migraine participants with and without visual aura.^11^

Participants were informed that the EMG electrodes measure an “eye response”. This wording was designed to avoid cueing the participant to the use of the EMG as an index of discomfort. While we presume that the EMG is an index of implicit discomfort, we did not assess what assumptions participants had about our measurement.

### The mechanism of signal amplification

While our results here and previously are consistent with a post-retinal amplification of ipRGC signals,^17^ the neural locus or physiologic mechanism of this amplification is not known. Although ipRGC signals appear to be amplified in both MwA and MwoA groups for explicit reports of discomfort, it is uncertain whether this effect is from the same mechanism in the two groups. A perhaps relevant finding is that MwA and MwoA groups differ in their cortical response to flickering light, with an enhanced response only seen in the MwA group.^29^

Given the dissociation of explicit and implicit discomfort measures in the presence and absence of visual aura, it seems likely that the physiologic mechanism of enhanced reflexive eye closure differs at some point from that of an enhanced conscious report of visual discomfort. There are several potential routes by which migraine might enhance a brainstem-mediated reflex. A decrease in habituation of the trigeminally-mediated blink reflex has been shown in migraine,^33, 34^ consistent with theories of a decrease in subcortical inhibitory processes in this condition.^35^ The calcitonin gene-related peptide (and other neuropeptides) are active in subcortical structures related to trigeminal discomfort.^36^ Alterations in these peptides is a feature of migraine and may account for the enhancement of brainstem reflexes related to discomfort.^36^

Trigeminal and retinal signals appear to interact and potentiate each other. In migraine, noxious trigeminal stimulation decreases visual discomfort thresholds,^5^ increases light-induced pain,^35^ and potentiates visual cortex activity in response to light.^37^ The reverse association is also found, as light decreases pain thresholds for trigeminal stimulation.^38^ In an fMRI study of a patient with transient photophobia from corneal irritation, visual stimulation was reported to produce activation of the trigeminal ganglion and trigeminal nucleus caudalis,^39^ in agreement with similar measurements in rodents.^16, 40, 41^ Trigeminal sensitization appears to facilitate photophobia in blepharospasm, a focal dystonia characterized by involuntary, repetitive eye closure.^42^ The ability of corneal irritation to induce light aversion is attenuated in mice lacking ipRGCs, suggesting a link between ipRGC signals and trigeminal nociception.^43^

An intriguing possibility is that melanopsin expression in trigeminally-innervated tissue could itself contribute to visual discomfort. Melanopsin has been found in the trigeminal ganglia of mice and humans,^44, 45^ and the cornea^45^ and iris of mice.^46^ In rodents with optic nerve lesions, bright light nonetheless potentiates the trigeminal blink reflex,^47^ and induces light aversion in a migraine-like state.^44^ In our study, however, extra-retinal melanopsin signaling seems unlikely to have played a substantial role, as the stimulus was transmitted through an artificial pupil into the pharmacologically dilated eye, minimizing the area of stimulated cornea and iris.

### Limitations

As we recruited participants with migraine from the community, not from a clinical setting, our studied population had episodic migraine with only moderate disability, and few participants reported using preventative medications. It is possible that the group effects we observe here would differ in a cohort of participants with chronic symptoms or a greater headache burden.

Our data analysis includes deviations from our pre-registered protocol. The most substantial of these is the use of a modified temporal window for the quantification of responses. While we feel that this modification is well justified, as it avoids a non-specific stimulus offset response, our statistical results are less robust when the pre-registered analysis approach is used. We observe in our data that an enhanced OO-EMG response does not appear to be a uniform feature of the MwA group. Relatedly, the distribution of responses across participants is greater for the MwA group as compared to the MwoA or headache-free control group. We do not have an explanation for this uneven distribution of effect size among the MwA participants.

## Conclusion

Our study shows light-induced, reflexive eye closure is enhanced in individuals with migraine with aura and interictal photophobia (MwA). These responses were evoked by melanopsin and cone stimulation, both in isolation and in combination, suggesting that ipRGCs are involved in the afferent arc of these reflexive responses. Our implicit measures of visual discomfort, OO-EMG activity and blink rate, were enhanced only in MwA, whereas in a prior study using an explicit rating of visual discomfort, we found enhanced responses in migraine regardless of a history of aura.^17^ These differences in outcomes between groups and measures suggests that ipRGC signals for discomfort are processed along multiple pathways, only some of which are altered in migraine in the absence of visual aura.

## Supplemental Materials

**Supplemental Figure 1.**
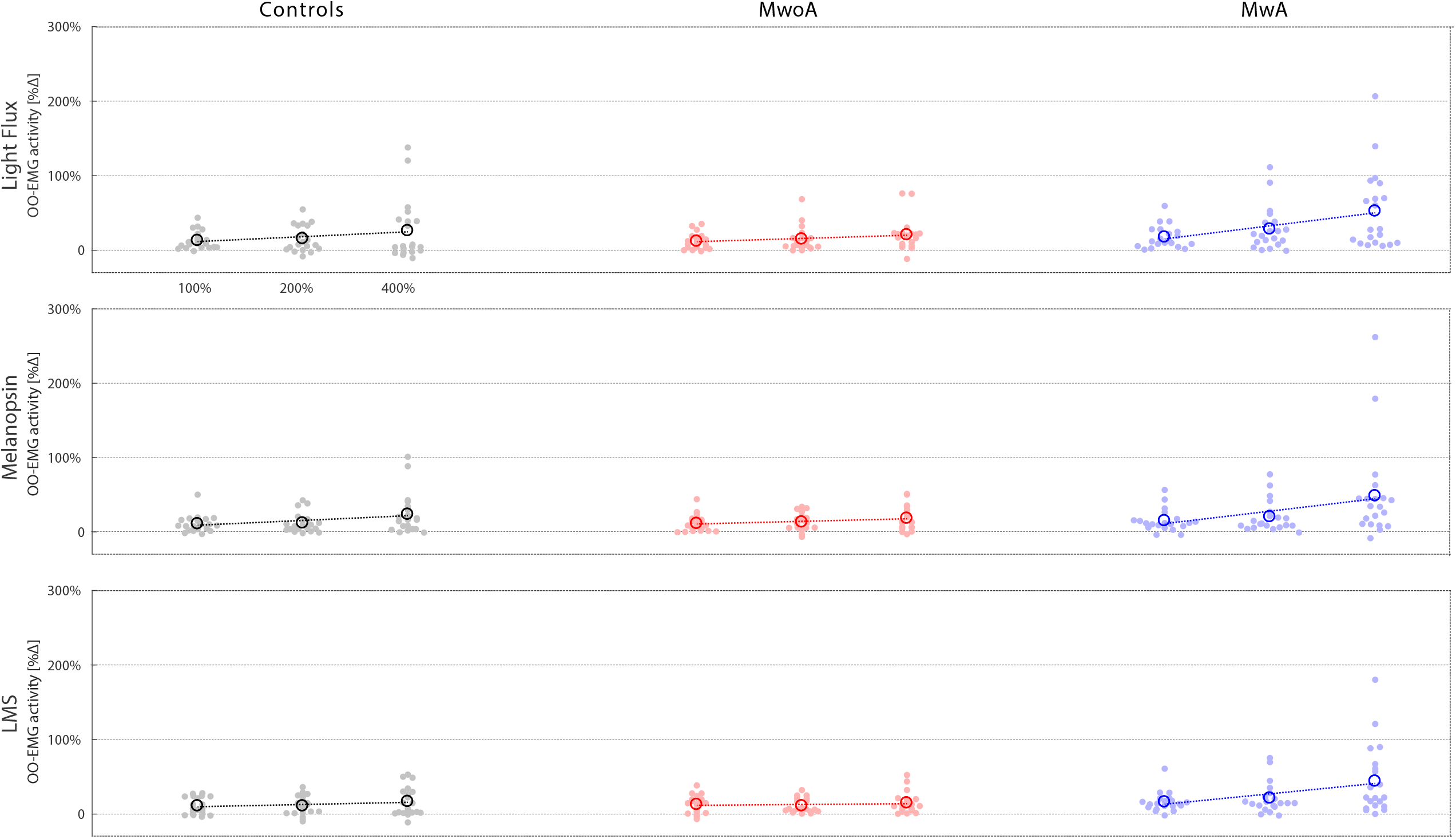
Orbicularis oculi electromyography (OO-EMG) activity by stimulus and group. Each row presents the OO-EMG activity evoked by stimuli that targeted a particular combination of photoreceptors, and each column contains the data from each individual group (n = 20 participants per group). The stimuli were presented at three different contrast levels (100, 200, and 400%), and these (log-spaced) values define the x-axis of each subplot. The mean (across trial) OO-EMG activity for a given stimulus and contrast is shown for each participant (filled circle), as is the mean OO-EMG activity across participants (open circle). The best fit line to the mean OO-EMG activity across participants as a function of log contrast is shown in each subplot. OO-EMG activity over time was quantified by calculating the standard deviation of voltage activity within a 500 msec window, sliding across time, which is expressed as percentage change [%Δ] relative to activity occurring prior to pulse onset. From these normalized time series, we calculated the average evoked response over a 4000 msec window starting 1 second after pulse onset.

**Supplemental Figure 2.**
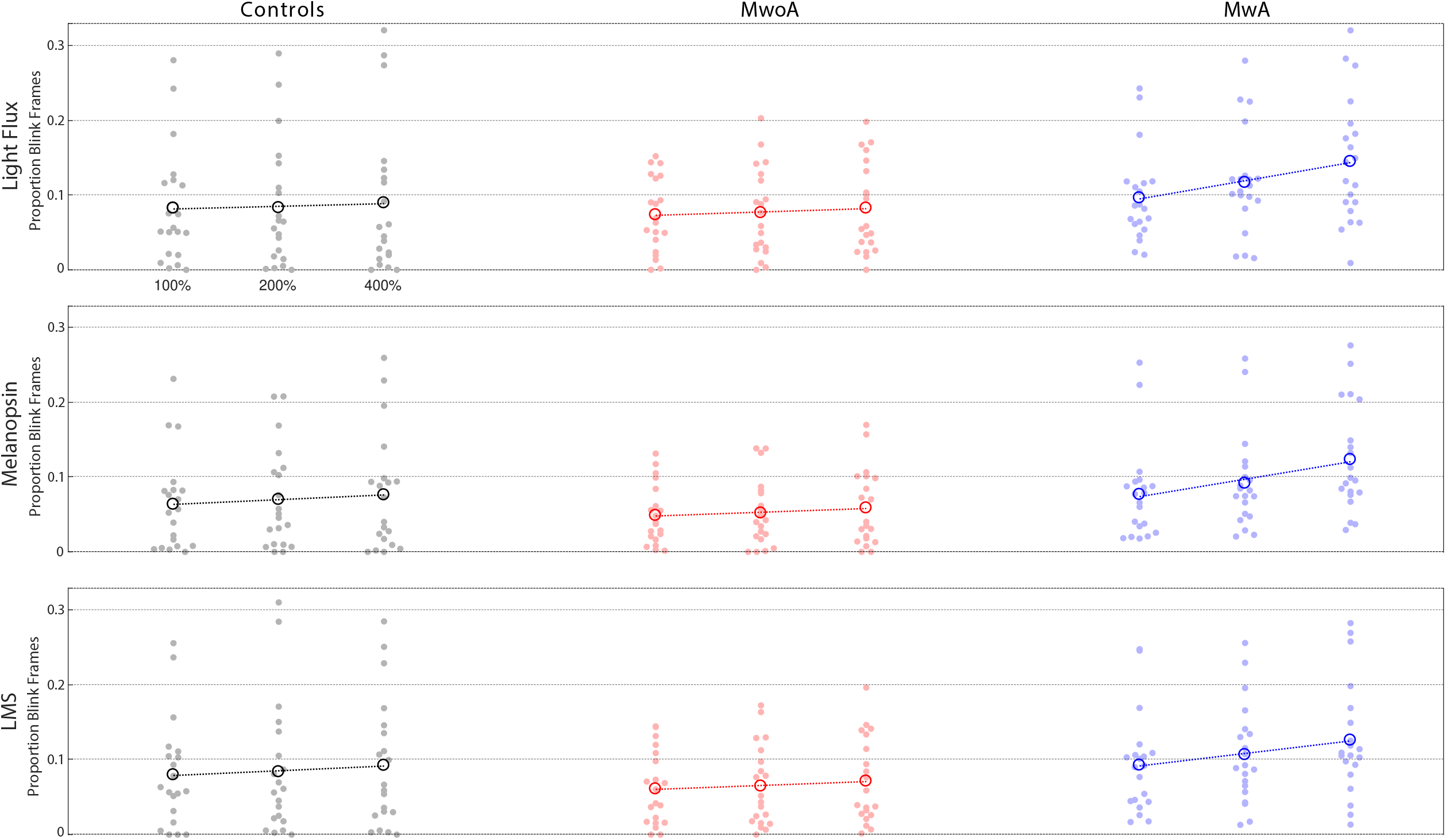
Blink activity by stimulus and group. Each row presents the blink activity evoked by stimuli that targeted a particular combination of photoreceptors, and each column contains the data from each individual group (n = 20 participants per group). The stimuli were presented at three different contrast levels (100, 200, and 400%), and these (log-spaced) values define the x-axis of each subplot. The mean (across trial) blink activity for a given stimulus and contrast is shown for each participant (filled circle), as is the mean blink activity across participants (open circle). The best fit line to the mean blink activity across participants as a function of log contrast is shown in each subplot. Blink activity is expressed as the proportion of frames of the video recording in which the eyelid covered the eye over a 4000 msec window starting 1 second after pulse onset.

**Supplemental Table 1.** Post-hoc analysis of the effect of contrast on orbicularis oculi EMG activity. We assessed the effect of contrast on OO-EMG activity within the 3400-msec analysis window by calculating the mean across-subject response at each log-spaced contrast level for a given group and stimulus type. Then we measured the slope for these three response values and obtained the 95% CI around that slope by bootstrapping across subjects.

| Group | Stimulus | Mean slope | 95% CI |
| --- | --- | --- | --- |
| *Controls* |  |  |  |
|  | Light Flux | 20.4 | 2.5 – 40.5 |
|  | Melanopsin | 7.8 | -3.4 – 19.9 |
|  | Cones | 7.5 | 1.1 – 13.8 |
| *MwoA* |  |  |  |
|  | Light Flux | 10.0 | -1.0 – 20.8 |
|  | Melanopsin | 7.8 | -0.6 – 16.7 |
|  | Cones | -0.9 | -6.8 – 4.6 |
| *MwA* |  |  |  |
|  | Light Flux | 58.7 | 30.4 - 92.4 |
|  | Melanopsin | 55.2 | 20.4 - 99.7 |
|  | Cones | 43.3 | 18.2 - 72.3 |

**Supplemental Table 2.** Post-hoc analyses of group effects by stimulus type in orbicularis oculi EMG activity at 400% contrast. We examined the effect of group and stimulus upon the mean OO-EMG activity using a 3-by-3 mixed effects ANOVA. This analysis was conducted over a 3400-msec window starting 0.3 second after pulse onset and ending 0.3 second before pulse offset, which was briefer than our pre-registered analysis. This smaller window was designed to avoid observed onset and offset responses. Independent variables included group (migraine with aura, without aura, and control) as a between subjects factor and photoreceptor target (melanopsin, cones, and light flux) as a within subjects factor. Post-hoc testing was conducted using the Tukey procedure. *p-value < 0.05.

| Stimulus | Group comparison | Mean difference | t-statistic | p-value |
| --- | --- | --- | --- | --- |
| *Light Flux* | MwA - Controls | 28.5 | 2.259 | 0.070 |
|  | MwA - MwoA | 34.6 | 2.736 | 0.022* |
|  | MwoA - Controls | -6.0 | -0.477 | 0.882 |
| *Melanopsin* | MwA - Controls | 31.6 | 2.391 | 0.052 |
|  | MwA - MwoA | 33.7 | 2.546 | 0.036* |
|  | MwoA - Controls | -2.1 | -0.156 | 0.987 |
| *Cones* | MwA - Controls | 28.8 | 3.107 | 0.008* |
|  | MwA - MwoA | 30.7 | 3.332 | 0.004* |
|  | MwoA - Controls | -2.0 | -0.215 | 0.975 |

**Supplemental Table 3.**
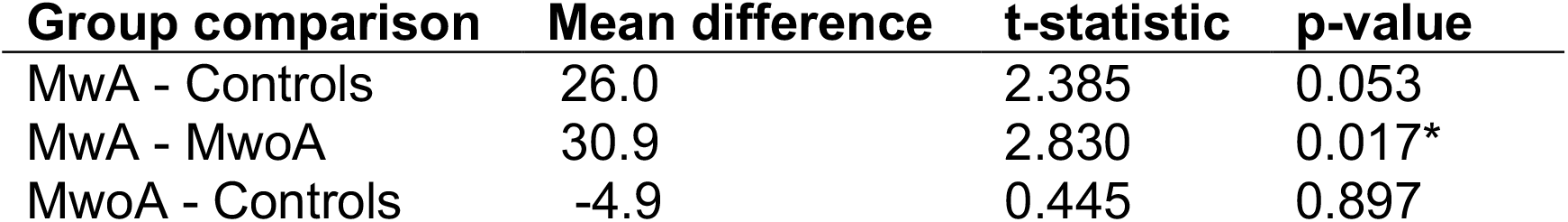
Post-hoc analyses of group effects in orbicularis oculi EMG activity at 400% contrast. We examined the effect of group upon the mean OO-EMG activity using a 3-by-3 mixed effects ANOVA. This analysis was conducted over a 4000-msec response window starting 1 second after pulse onset as per our preregistration document. Measured EMG activity across stimulus types (melanopsin, cones, and light flux) were averaged within subjects. The independent variables were by group (MwA, MwoA, and control) as a between subjects factor. Post-hoc testing was conducted using the Tukey procedure. *p-value < 0.05.

| **Group comparison** | **Mean difference** | **t-statistic** | **p-value** |
| --- | --- | --- | --- |
| MwA - Controls | 26.0 | 2.385 | 0.053 |
| MwA - MwoA | 30.9 | 2.830 | 0.017* |
| MwoA - Controls | -4.9 | 0.445 | 0.897 |

**Supplemental Table 4.**
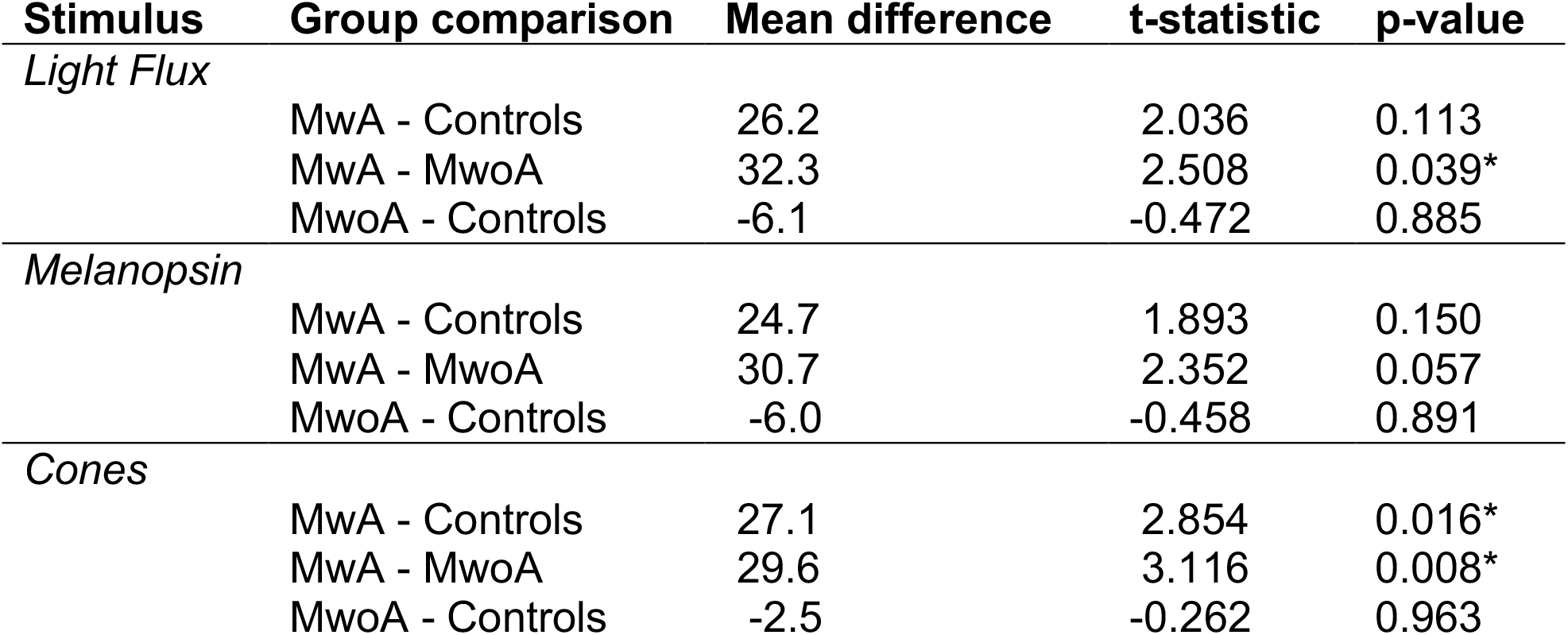
Post-hoc analyses of group effects by stimulus type in orbicularis oculi EMG activity at 400% contrast. We examined the effect of group and stimulus upon the mean OO-EMG activity using a 3-by-3 mixed effects ANOVA. This analysis was conducted over a 4000-msec response window starting 1 second after pulse onset as per our preregistration document. Independent variables included group (migraine with aura, without aura, and control) as a between subjects factor and photoreceptor target (melanopsin, cones, and light flux) as a within subjects factor. Post-hoc testing was conducted using the Tukey procedure. *p-value < 0.05.

| Stimulus | Group comparison | Mean difference | t-statistic | p-value |
| --- | --- | --- | --- | --- |
| *Light Flux* | MwA - Controls | 26.2 | 2.036 | 0.113 |
|  | MwA - MwoA | 32.3 | 2.508 | 0.039* |
|  | MwoA - Controls | -6.1 | -0.472 | 0.885 |
| *Melanopsin* | MwA - Controls | 24.7 | 1.893 | 0.150 |
|  | MwA - MwoA | 30.7 | 2.352 | 0.057 |
|  | MwoA - Controls | -6.0 | -0.458 | 0.891 |
| *Cones* | MwA - Controls | 27.1 | 2.854 | 0.016* |
|  | MwA - MwoA | 29.6 | 3.116 | 0.008* |
|  | MwoA - Controls | -2.5 | -0.262 | 0.963 |

**Supplemental Table 5.** Post-hoc analysis of the effect of contrast on blink activity. We assessed the effect of contrast on proportion of blink frames within the 3400-msec analysis window by calculating the mean across-subject response at each log-spaced contrast level for a given group and stimulus type. Then we measured the slope for these three response values and obtained the 95% CI around that slope by bootstrapping across subjects.

| Group | Stimulus | Mean slope | 95% CI |
| --- | --- | --- | --- |
| *Controls* |  |  |  |
|  | Light Flux | -0.0064 | -0.0220 – 0.0102 |
|  | Melanopsin | -0.0068 | -0.0238 – 0.0105 |
|  | Cones | -0.0043 | -0.0125 – 0.0042 |
| *MwoA* |  |  |  |
|  | Light Flux | 0.0145 | -0.0022 – 0.0312 |
|  | Melanopsin | 0.0018 | -0.0087 – 0.0116 |
|  | Cones | 0.0043 | -0.0023 – 0.0113 |
| *MwA* |  |  |  |
|  | Light Flux | 0.0726 | 0.0393 – 0.1106 |
|  | Melanopsin | 0.0549 | 0.0283 – 0.0824 |
|  | Cones | 0.0478 | 0.0166 – 0.0829 |

**Supplemental Table 6.** Post-hoc analyses of group effects by stimulus type in blink activity at 400% contrast. We examined the effect of group and stimulus upon the mean proportion of blink frames using a 3-by-3 mixed effects ANOVA. This analysis was conducted over a 3400-msec window starting 0.3 second after pulse onset and ending 0.3 second before pulse offset, which was briefer than our pre-registered analysis. This smaller window was designed to avoid observed onset and offset responses. Independent variables included group (migraine with aura, without aura, and control) as a between subjects factor and photoreceptor target (melanopsin, cones, and light flux) as a within subjects factor. Post-hoc testing was conducted using the Tukey procedure. *p-value < 0.05.

| Stimulus | Group comparison | Mean difference | t-statistic | p-value |
| --- | --- | --- | --- | --- |
| *Light Flux* | MwA - Controls | 0.0689 | 3.257 | 0.005* |
|  | MwA - MwoA | 0.0561 | 2.655 | 0.027* |
|  | MwoA - Controls | 0.0127 | 0.602 | 0.820 |
| *Melanopsin* | MwA - Controls | 0.0510 | 3.048 | 0.010* |
|  | MwA - MwoA | 0.0592 | 3.540 | 0.002* |
|  | MwoA - Controls | -0.0082 | -0.492 | 0.875 |
| *Cones* | MwA - Controls | 0.0489 | 2.683 | 0.025* |
|  | MwA - MwoA | 0.0485 | 2.663 | 0.027* |
|  | MwoA - Controls | 0.0004 | 0.020 | 0.999 |

**Supplemental Table 7.**
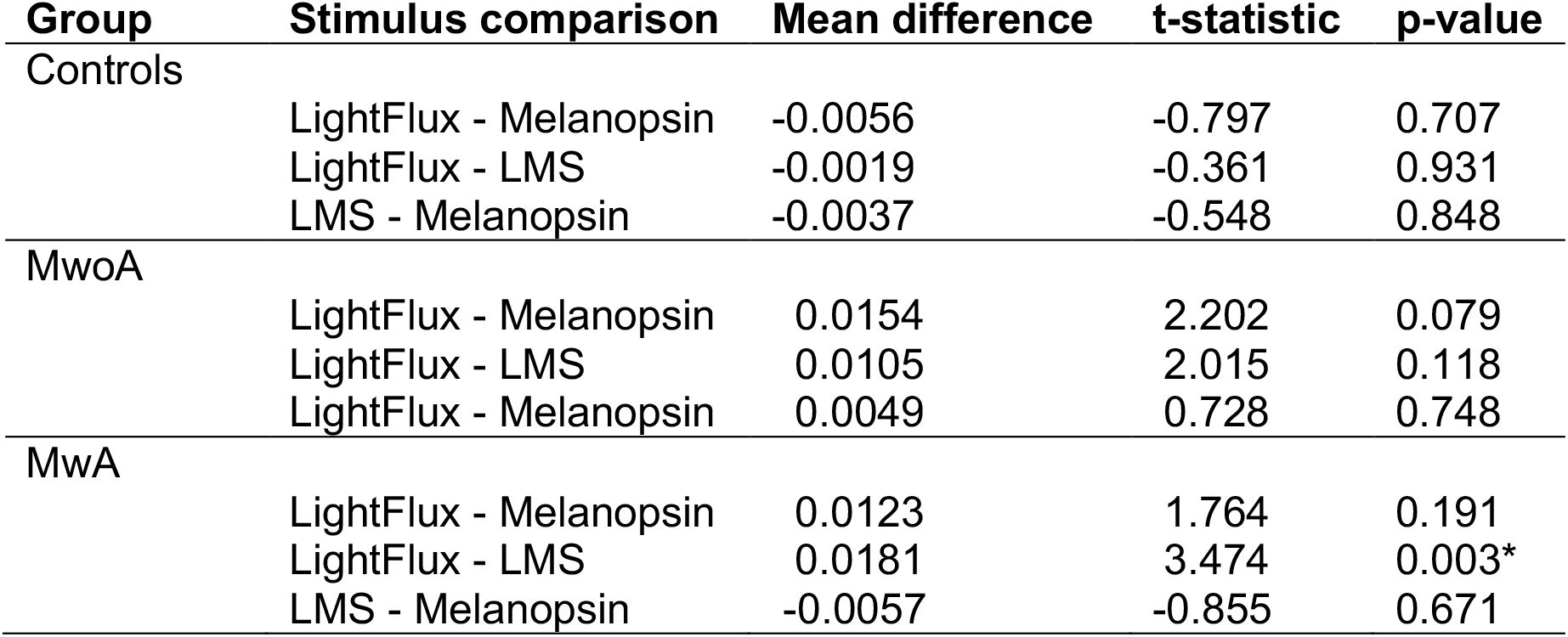
Post-hoc analyses of stimulus type effects by group in blink activity at 400% contrast. We examined the effect of group and stimulus upon the mean proportion of blink frames using a 3-by-3 mixed effects ANOVA. This analysis was conducted over a 3400-msec window starting 0.3 second after pulse onset and ending 0.3 second before pulse offset, which was briefer than our pre-registered analysis. This smaller window was designed to avoid observed onset and offset responses. Independent variables included photoreceptor target (melanopsin, cones, and light flux) as a between subjects factor and group (migraine with aura, without aura, and control) as a within subjects factor. Post-hoc testing was conducted using the Tukey procedure. *p-value < 0.05.

| Group | Stimulus comparison | Mean difference | t-statistic | p-value |
| --- | --- | --- | --- | --- |
| Controls | LightFlux - Melanopsin | -0.0056 | -0.797 | 0.707 |
|  | LightFlux - LMS | -0.0019 | -0.361 | 0.931 |
|  | LMS - Melanopsin | -0.0037 | -0.548 | 0.848 |
| MwoA | LightFlux - Melanopsin | 0.0154 | 2.202 | 0.079 |
|  | LightFlux - LMS | 0.0105 | 2.015 | 0.118 |
|  | LightFlux - Melanopsin | 0.0049 | 0.728 | 0.748 |
| MwA | LightFlux - Melanopsin | 0.0123 | 1.764 | 0.191 |
|  | LightFlux - LMS | 0.0181 | 3.474 | 0.003* |
|  | LMS - Melanopsin | -0.0057 | -0.855 | 0.671 |

**Supplemental Table 8.** Post-hoc analyses of group effects in blink activity at 400% contrast. We examined the effect of group upon the mean proportion of blink frames using a 3-by-3 mixed effects ANOVA. This analysis was conducted over a 4000-msec response window starting 1 second after pulse onset as per our preregistration document. Responses across stimulus types (melanopsin, cones, and light flux) were averaged within subjects. The independent variables were by group (MwA, MwoA, and control) as a between subjects factor. Post-hoc testing was conducted using the Tukey procedure. *p-value < 0.05.

| **Group comparison** | **Mean difference** | **t-statistic** | **p-value** |
| --- | --- | --- | --- |
| MwA - Controls | 0.0459 | 1.958 | 0.132 |
| MwA - MwoA | 0.0614 | 2.614 | 0.030* |
| MwoA - Controls | -0.0154 | 0.656 | 0.790 |

**Supplemental Table 9.** Post-hoc analyses of group effects by stimulus type in blink activity at 400% contrast. We examined the effect of group and stimulus upon the mean proportion of blink frames using a 3-by-3 mixed effects ANOVA. This analysis was conducted over a 4000-msec response window starting 1 second after pulse onset as per our preregistration document. Independent variables included group (migraine with aura, without aura, and control) as a between subjects factor and photoreceptor target (melanopsin, cones, and light flux) as a within subjects factor. Post-hoc testing was conducted using the Tukey procedure. *p-value < 0.05.

| Stimulus | Group comparison | Mean difference | t-statistic | p-value |
| --- | --- | --- | --- | --- |
| *Light Flux* | MwA - Controls | 0.0559 | 2.109 | 0.097 |
|  | MwA - MwoA | 0.0630 | 2.378 | 0.053 |
|  | MwoA - Controls | -0.0072 | -0.270 | 0.961 |
| *Melanopsin* | MwA - Controls | 0.0479 | 2.193 | 0.081 |
|  | MwA - MwoA | 0.0657 | 3.007 | 0.011* |
|  | MwoA - Controls | -0.0178 | -0.814 | 0.696 |
| *Cones* | MwA - Controls | 0.0341 | 1.434 | 0.330 |
|  | MwA - MwoA | 0.0554 | 2.329 | 0.060 |
|  | MwoA - Controls | -0.0213 | -0.894 | 0.646 |

**Supplemental Table 10.** Post-hoc analyses of stimulus type effects by group in blink activity at 400% contrast. We examined the effect of group and stimulus upon the mean proportion of blink frames using a 3-by-3 mixed effects ANOVA. This analysis was conducted over a 4000-msec response window starting 1 second after pulse onset as per our preregistration document. Independent variables included photoreceptor target (melanopsin, cones, and light flux) as a between subjects factor and group (migraine with aura, without aura, and control) as a within subjects factor. Post-hoc testing was conducted using the Tukey procedure. *p-value < 0.05.

| Group | Stimulus comparison | Mean difference | t-statistic | p-value |
| --- | --- | --- | --- | --- |
| Controls | LightFlux - Melanopsin | 0.0133 | 1.770 | 0.189 |
|  | LightFlux - LMS | -0.0030 | -0.514 | 0.865 |
|  | LMS - Melanopsin | 0.0163 | 2.403 | 0.050 |
| MwoA | LightFlux - Melanopsin | 0.0239 | 3.187 | 0.007* |
|  | LightFlux - LMS | 0.0111 | 1.889 | 0.151 |
|  | LightFlux - Melanopsin | 0.0128 | 1.889 | 0.151 |
| MwA | LightFlux - Melanopsin | 0.0213 | 2.835 | 0.017* |
|  | LightFlux - LMS | 0.0187 | 3.189 | 0.006* |
|  | LMS - Melanopsin | 0.0025 | 0.373 | 0.926 |

